# Food hardness and stone tool weight in wild primate nut-cracking

**DOI:** 10.1101/267542

**Authors:** Michael Haslam

## Abstract

This study presents data on average stone tool weights and the hardness of foods processed by the three known stone-tool-using primate species: Burmese long-tailed macaques (*Macaca fascicularis aurea*), bearded capuchins (*Sapajus libidinosus*) and Western chimpanzees (*Pan troglodytes verus*). Each of these primates uses stone hammers to crack open nuts in the wild, making them suitable for inter-species behavioural comparison. This work draws on published results to identify a distinct difference in the tool weight/food hardness curve between chimpanzees and the two monkey taxa, with the latter reaching an asymptote in mean tool weight of just over 1 kg regardless of increasing food hardness. In contrast, chimpanzees rapidly increase their tool weight in response to increasing hardness, selecting average masses over 5 kg to process the hardest nuts. Species overlap in their preference for tools of 0.8-1 kg for opening foods of hardness 2-3 kN, suggesting that this conjunction may represent a primate stone-tool-use optimum.

## Introduction

Stone tools have been an essential component of the hominin (human lineage) toolkit for millions of years (Harmand et al. 2015). Until the late twentieth century, however, our primary reference for the use of such tools was other hominins, including modern humans (Reybrouck 2012). With the discovery that some members of the West African chimpanzee sub-species (*Pan troglodytes verus*) used stones to break open nuts in the wild, direct comparison with the behaviour of these living apes became feasible (Joulian 1996; Marchant and McGrew 2005; Whiten et al. 2009; Boesch 2012). Subsequently, bearded capuchin monkeys (*Sapajus libidinosus*) in Brazil (Ottoni and Izar 2008; Proffitt et al. 2016) and Burmese long-tailed macaques (*Macaca fascicularis aurea*) in Thailand (Gumert and Malaivijitnond 2012; Haslam et al. 2016b) were also found to use stone pounding tools in their natural habitats. However, despite the fact that the two monkeys are equally as phylogenetically related to modern chimpanzees as they are to modern humans, hominin comparisons still dominate discussions of this behaviour (Visalberghi et al. 2015; Falótico et al. 2017).

Here, I compare the one type of stone tool use known to be shared by these three nonhuman primates (hereafter, primates): nut-cracking. Specifically, I examine how the hardness of a processed food influences the selected weight of stone pounding tools. These two variables have been shown to be significantly correlated in single-species studies (Boesch and Boesch 1983; Spagnoletti et al. 2011; Gumert and Malaivijitnond 2013), however they have not previously been assessed in an inter-species study.

In-depth coverage of the sites and types of wild stone tool use in these primates may be found elsewhere (Gumert et al. 2009; Matsuzawa 2011; Boesch 2012; Visalberghi and Fragaszy 2013; Falótico and Ottoni 2016). In summary, bearded capuchins commonly use handheld stones to pound open encased nuts, including palm nuts (*Attalea, Astrocarpum* and *Orbignya*) and cashews (*Anacardium* sp.) (Visalberghi and Fragaszy 2013; Mendes et al. 2015; Luncz et al. 2016b). Burmese long-tailed macaques primarily use handheld stones to process intertidal oysters and gastropods (Gumert and Malaivijitnond 2012), however they have also been observed to pound open nuts of sea almonds (*Terminalia catappa*) (Falótico et al. 2017) and oil palms (*Elaeis guineensis*) (Luncz et al. 2017). Chimpanzees stone tool use is focused on cracking a variety of nuts and fruits (including *Panda*, *Coula*, *Parinari* and *Elaeis*) (McGrew 1992). All three primates select and transport stones to tool-use sites, for use as hammers (Visalberghi et al. 2013; Haslam et al. 2016b; Luncz et al. 2016a) and anvils (Sakura and Matsuzawa 1991; Haslam et al. 2016a).

## Methods

I collated available published data on wild primate activities that involve use of a handheld hammerstone to break open encased food, where both tool weights and the food hardness are available. With one exception (see below), I only included behaviour recorded in natural settings, rather than from captive or experimental situations (Matsuzawa 1994; Visalberghi et al. 2009; Luncz et al. 2016b), in order to avoid biases that humans may introduce to primate performance (Haslam 2013). Data were drawn from the literature, except for the average weight of tools (n=348) used by wild macaques to process oil palm nuts on Koh Yao Noi, Thailand. The latter data were recorded during surveys in October 2016 as part of the ERC-funded Primate Archaeology project.

To ensure comparability, I included only measures of food hardness expressed as fracture force (in kN). This variable is the one most relevant to primate stone pounding behaviour, as opposed to food toughness, for example (Berthaume 2016). I used average tool weights, combined for both sexes, because researchers rarely provided raw data that would permit more precise comparisons. Tool data were only included when the report indicated that a single species of nut was processed with those tools. For the purposes of this initial report, I have not taken tool hardness or material into account, although I acknowledge that these factors will affect the effectiveness of a stone hammer of a given weight.

Tool weights from the capuchin Fazenda Boa Vista (FBV) site have been reported in terms of their use on ‘high resistance’ and ‘low resistance’ nuts (Spagnoletti et al. 2011), and I have followed that procedure here, leading to identical weights for tools used to process each of the nut species that fall into these categories (see Table 1). Chimpanzee tools from the Taï Forest have been published in weight categories (Boesch and Boesch 1984; Luncz et al. 2016a), and an average value was therefore calculated by assigning each category a weight in the centre of its range (e.g., the category ‘1-2.9 kg’ was assigned a value of 2 kg) (Boesch and Boesch 1984). The macaque data include the two nut species processed with stone tools; however, I also included two gastropod species to reflect the fact that these monkeys predominantly use stones on shellfish. The gastropods include relatively easily-opened (*Nerita* sp.) and hard-shelled (*Thais* sp.) prey. The macaque shell-cracking data are the exception to my excluding experimentally provided stones (Gumert and Malaivijitnond 2013), as mollusc-specific tool weights are not otherwise available.

## Results and Discussion

Results are presented in Table 1 and Fig. 1. The results replicate previous findings that tool selection, in terms of average weight, is positively correlated with food hardness for all three non-human primate stone-tool-using species. However, the inter-species comparison reveals that this is not a linear process. Instead, each of the primates asymptotically approaches a maximum average tool weight, where increasing food hardness results in successively smaller increases in selected tool weight.

**Fig. 1.**
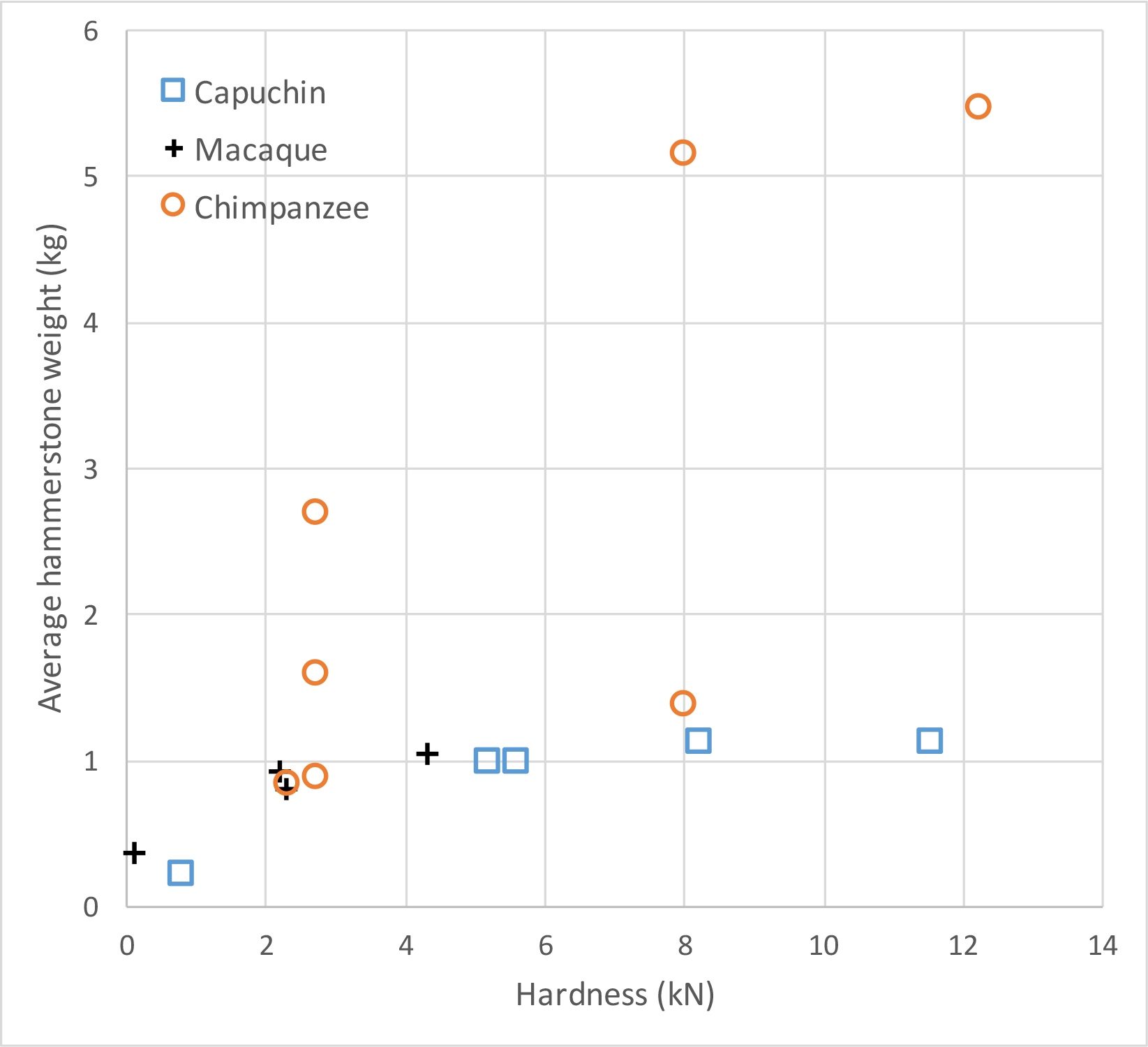
Food hardness (kN) and associated average hammerstone weight (kg) for wild non-human primates

**Table 1.**
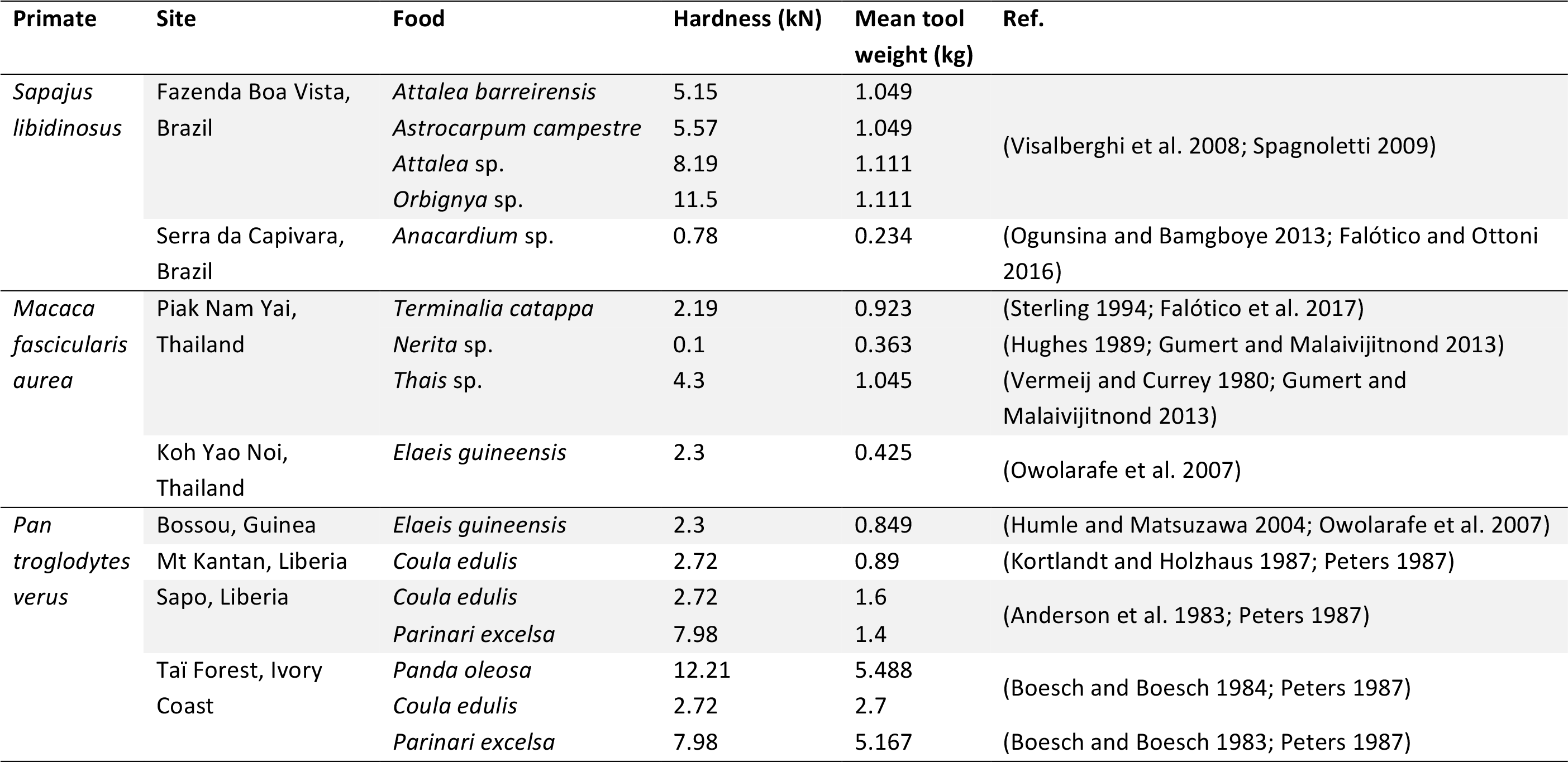
Food hardness and associated stone tool weights for wild non-human primates

The two monkey species follow approximately the same shallow curve, with each using lighter tools to process softer materials (*Nerita* for macaques, cashews for capuchins), and reaching a maximum average of just over a kilogram for tools used on the hardest capuchin palm nuts and macaque molluscs. Heavier stones suitable for use as hammers are available, albeit not abundant, at all the well-studied monkey tool-use sites (MH, pers. obs.), which suggests that stone availability is not solely dictating this asymptote. Instead, the limiting factor may be body size and associated strength in manipulating stones. Supporting this suggestion, female capuchins at FBV used significantly lighter hammerstones when cracking low resistance as opposed to high resistance nuts, whereas the larger males showed no such distinction (Spagnoletti et al. 2011). Similarly, male macaques on Piak Nam Yai used heavier stones significantly more often than did the smaller females (Gumert et al. 2011). However, long-tailed macaques typically weigh 4-7 kg (Hamada et al. 2008) and bearded capuchins 2-4 kg (Fragaszy et al. 2016), and if body size is an important determinant of tool size (Visalberghi et al. 2015), then we may expect the macaque curve to level out above that of the capuchins. Testing this hypothesis will require finding wild *M.f. aurea* populations that have naturally developed stone tool use on harder foods than those currently processed.

The chimpanzee curve rises much faster than that of either of the monkeys, beginning to level out only above 5 kg tool weight, and not yet reaching a clear asymptote. Given that foods of similar hardness are cracked by both chimpanzees and capuchins (for example, *Panda* and *Orbignya*), the stark discrepancy between these two taxa is again most likely explained by differences in strength derived from body weight. Adult Western chimpanzees can weigh approximately 45-55 kg, and the wild capuchins therefore need to use stones that weigh a significantly higher proportion of their body mass to break open highly resistant nuts (Visalberghi et al. 2015). Complicating this picture slightly, one chimpanzee data point–for processing *Parinari* at Sapo in Liberia–falls closer to the monkey curves, indicating that there is likely more variation to be uncovered as additional food-specific tool choices are recorded under natural conditions. In any case, the data suggest that wild stone-tool-using chimpanzees are capable of cracking open items of higher maximum hardness than has been recorded to date. Whether foods or other objects of such hardness exist in their habitats, and whether the chimpanzees would gain from breaking them open, requires further study.

The dominance of average weights in the published literature, and relied upon here, obscures extremes and may bias the current study in unknown ways. For example, chimpanzees in the Taï Forest have been reported to use tools up to 24 kg (Boesch and Boesch 1983), and capuchins at Fazenda Boa Vista were observed to occasionally use hammers up to 3 kg (Spagnoletti et al. 2011). Nevertheless, the use of mean weights has the benefit of focusing on common behaviour, not just that of the strongest–typically male–individuals. Reporting of more complete datasets in the future will aid in parsing the necessarily broad conclusions drawn in this initial investigation, as will further characterization of the hardness of foods pounded by non-human primates.

The emphasis on averages does permits direct comparisons across all three nut-cracking non-human primates, potentially identifying cross-taxa preferences. For example, the monkey and chimpanzee curves meet at a collection of data points between 2-3 kN and 0.8-1 kg. On present evidence, this combination of food hardness and tool weight may therefore record something of an inter-species stone tool use optimum for primates.

Although comparison between extant primate and extinct hominin percussion was not the aim of this study, the inclusion of hominin percussive hammerstone data from sites where pounding activities and consumed foods can be inferred (Arroyo and de la Torre 2016) would act as a test of this suggestion. Finally, if body mass is indeed a driver of tool weight, then hominin stone tool selection would be expected to more closely follow the chimpanzee curve rather than those of the monkeys. Similarly, if additional extant stone-tool-using primates are discovered, their percussive preferences will add valuable extra information to the comparative data against which past primate and hominin behavioral reconstructions can be assessed (Haslam et al. 2017).

## Acknowledgements

I thank the members of the Primate Archaeology research network for their advice, and Lydia Luncz, Magda Svensson and Michael Gumert for their assistance in Thailand. This work was funded by a European Research Council Starting Grant (no. 283959) awarded to MH.

